# Deficient stereopsis in the normal population revisited: why current clinical stereo tests may not be adequate

**DOI:** 10.1101/585976

**Authors:** Robert F Hess, Rebecca Dillon, Rifeng Ding, Jiawei Zhou

## Abstract

**Significance statement:** Applied applications for occupational screening, clinical tests should assess sensitivity to the sign as well as the magnitude of disparity.

**Purpose:** To determine why the high incidence of stereo anomaly found using laboratory tests with polarity-based increment judgements (i.e., depth sign) is not reflected in clinical measurements that involve single-polarity incremental judgements (i.e., depth magnitude).

**Methods:** An iPod-based measurement that involved the detection of an oriented shape defined by a single polarity-depth increment within a random dot display was used. A staircase procedure was used to gather sufficient trials to derive a meaningful measure of variance for the measurement of stereopsis over a large disparity range. Forty-five adults with normal binocular vision (20 - 65 years old) and normal or corrected-to-normal (0 logMAR or better) monocular vision participated in this study.

**Results:** Observers’ stereo acuities ranged between 10 and 100 arc seconds, and were normally distributed on a log scale (*p* = 0.90, 2-tailed Shapiro-Wilk test). The present results using a single polarity depth increment task (i.e., depth magnitude) show a similar distribution to those using a similar task using the Randot preschool stereo test on individuals between the ages of 19-35 using either the 4-book test (n = 33) or the 3-book test (n = 40), but very different results when the iPod test involved a polarity-based increment judgement (i.e., depth sign).

**Conclusions:** The present clinical stereo tests are based on magnitude judgements and are unable to detect the high percentage of stereo anomalous individuals in the normal population revealed using depth sign judgements.

---

Recently, we developed a convenient method ^1^ for measuring stereopsis for the clinic using an iPod, which involves depth discrimination of clouds of random dots; the subject was presented with two circular stimuli in depth side-by-side, one was in front of the screen (crossed disparity), the other was behind the screen (uncrossed disparity); the subject was asked to detect which of the two stimuli was behind the screen (i.e., a depth sign judgement). Disparity as varied using a staircase procedure until a threshold was reached. One key finding was that as many as 30% of people with otherwise normal vision were over 10 times worse than their fellows. While this might be surprising to clinicians because the present clinical tests do not reflect this^2^, it is not surprising to vision scientists, as there have been a number of laboratory studies that have suggested this ^1, 3–9^. Although the reason for this stereo deficiency is presently unclear, Richards 4, 5 originally suggested that it may reflect a loss of a subset of disparity-sensitive neurons that process just crossed or just uncrossed disparities. One plausible reason why the current clinical tests ^2^ have failed to detect this is because they rely exclusively on detection of depth magnitude (i.e., a single polarity increment judgement) rather than depth sign. To test this, we have developed a different version of our previously published iPod clinical test^1^ that involves a detection of depths of fixed polarity (i.e., single polarity depth increment, as the majority of current clinical approaches do) rather than one that is dependent of depth polarity ^1^. We show that the high prevalence of stereo-deficiency within the normal population can only be revealed using a task that is dependent on depth polarity.

## Methods

### Observers

Forty-five adults (20 - 65 years old) with normal or corrected-to-normal (0 logMAR or better) monocular vision and no history of binocular dysfunction participated. Written consent was obtained prior to the study, which was approved by the Institutional Review Board of McGill University. The described research adhered to the tenets of the Declaration of Helsinki. Except the authors, all subjects were naive to the purpose of this study. The recruitment process and population demographics were essentially identical to that used previously^1^.

### Apparatus and stimulus

The measurement was conducted by a Mac iPod (Model No: A1367) running Stereogram Test app, an in-house software for IOS devices that featured a 326 ppi (pixels per inch) Retina Display. The app software was written in Objective-C using IOS software development kit combined with OpenGL ES 2.0. Observers viewed the stimuli dichoptically through red-green anaglyph glasses at a viewing distance of 50 cm in a normal light environment.

The stimulus was a pacman defined by static random-dots in depth on a random dot background, as shown in Figure 1A. The pacman was Gaussian-windowed to blend the edge to the background and to reduce monocular cues. The pacman contained randomly positioned red and green dots, which had a certain offset to generate depth perception (i.e., disparity). Overlapped red and green dots (or overlapped parts of dots, determined by the size of the dots) were blended into an orange color by using the blending functions provided by OpenGL ES to provide sub-pixel resolution.

**Figure 1.**
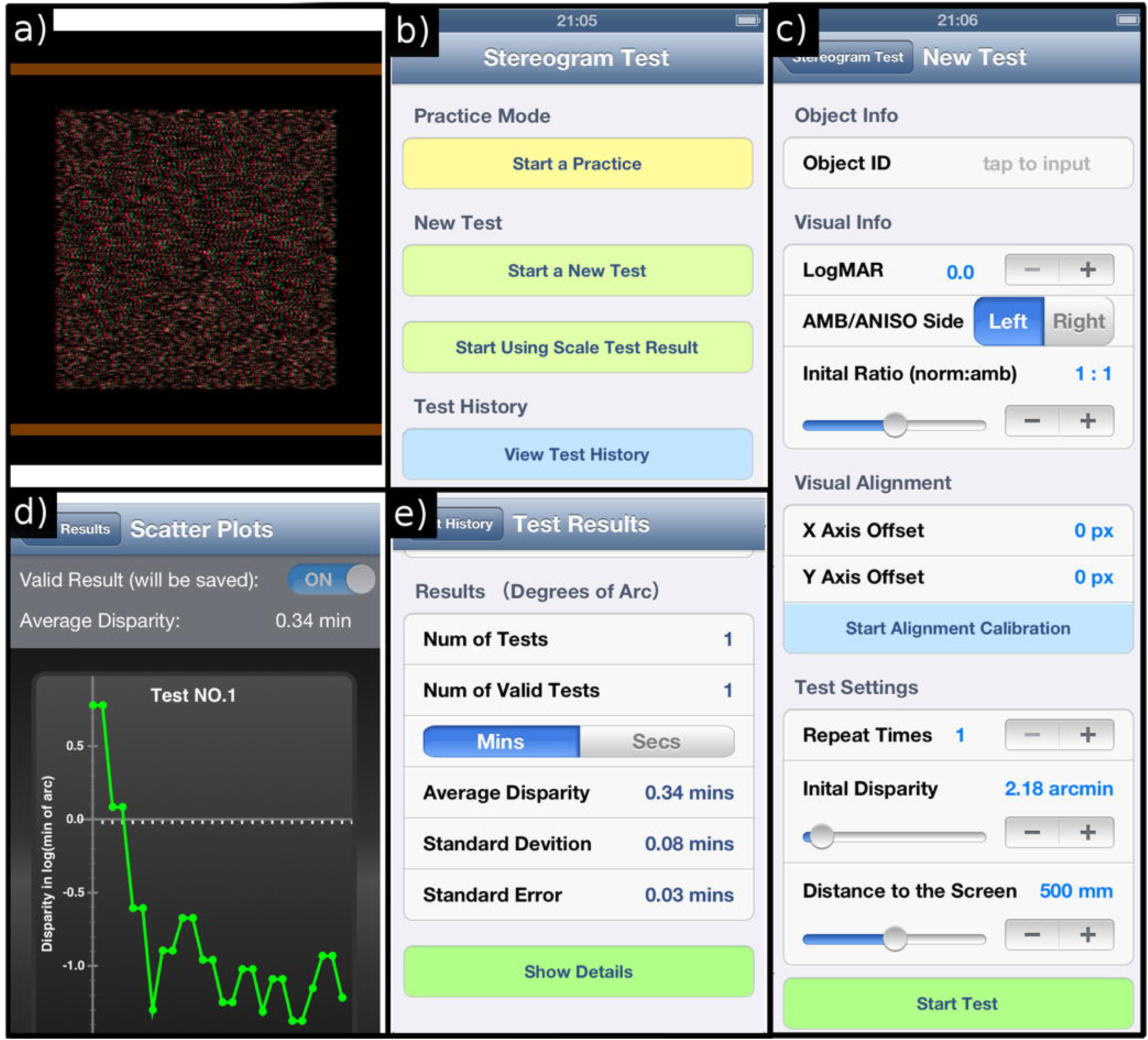
An illustration of the Stereogram Test App. (a) The random-dot stimuli; (b) a screenshot of the menus in this App; (c) the test configurations in the test-no aniso scaling or alignment adjustment was required in this study; (d) a plot of results after finishing the test; (e) a summary of the test results.

The offsets between red and green dots were equal thus the pacman was perceived as in front of the screen plane defined by the background random dots. In each trial, observers’ task was swipe the screen in the direction the pacman pointed; left, right, up or down. There was no time limit for responding, as the next trial came immediately after the observers’ response.

### Procedure

Observers were asked to finish a 10-trial practice before the test. After that, a 5-min test session, incorporating two separate runs, was used to estimate individuals’ stereo thresholds. Each run was driven by a staircase procedure in which, the initial offset between red and green dots was adjustable (could set up to 40 pixels, i.e., corresponding to a stereo acuity of 21.79 arc minutes) and was controlled by a 2-down/1-up staircase procedure thereafter. The initial step size was 50% (relative rate), which changed after the first reversal to 10% in all following trials. Since all of our participants were adults with normal vision, we set the maximal offset between red and green dots to 40 pixels to ensure that it was well within *D*_*max*_ (which was around 50-70 arc min) ^10^. The staircase was terminated at the fourth reversal point. The stereo threshold and its standard error were then calculated based on the last three reversals averaged across the two test runs (i.e., six reversals in total).

### Test Configuration

As is shown in Figure 1c, the following configurable parameters were provided in the Stereogram Test app:

1. Visual acuity of the worse eye;
2. AMB/ANISO Side and Initial Ratio: In case of spectacle-corrected anisometropia (with or without amblyopia), an image size-scaling feature was implemented to account for any aniseikonia. A second program was incorporated within this app to test the degree of aniseikonia; the size of the pixels could be scaled in front of the more emmetropic eye during the test to eliminate any potential impact of aniseikonia. This was a part of the original clinical iPod test^1^ and was designed for testing anisometropic amblyopes or subjects with a high degree of anisometropia. It was not utilized in this study because none of our subjects had a high degree of anisometropia (i.e., > 3 dioptres). Thus, the initial ratio was set to 1:1.
3. Visual Alignment: In case of ocular misalignment (e.g., strabismus), an alignment calibration feature was implemented to allow fusion of the two eyes’ images. During the alignment, two half-cross (one in red and the other one in green) were dichoptically presented to the two eyes. Observers were asked to align the two half-cross into a perfect whole cross. The degree to which the alignment was stable from run to run can be then assessed, as the alignment offset is provided to the examiner. This was part of the original iPod clinical stereo test^1^ and designed for strabismic patients. It was not utilized in this study because none of our participants had an ocular misalignment. Thus, x and y offsets were set to 0 px.
4. Repeat times: Participants were allowed to repeat individual test runs in a test session if the staircase results were clearly of an anomalous form indicative of a poor determination as the result of, for example, an early response mistake (finger error).

### Result Transformation

As is shown in Figure 1d, upon completion of the test, a plot of disparity as a function of trial number is provided for each test run. The disparity was recorded in pixels during the measurement and was converted into minute of arc by using the following equation:

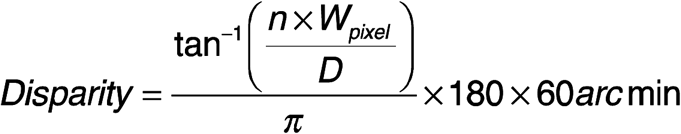

where *n* is the offsets between the red and green dots (in pixels), *W*_*pixel*_ is the physical width of a pixel on the display and *D* is the distance between the subject’s eyes and device’s display. In our study, *W*_*pixel*_ was 0.0792 mm and *D* was 500 mm.

## Results

Stereo acuity results for 45 normal observers are plotted in Figure 2a. The results using a single-polarity increment detection version of the test (Pacman) are normally distributed on this log plot between 10 and 100 arc seconds (*P* = .90, 2-tailed Shapiro-Wilk test). The present results show a similar distribution (on this log axis) to those of a previous study using the Randot preschool stereo test, which also involves detection of the shape defined by a single polarity depth increment, on individuals between the ages of 19-35 using either the 4-book test (n = 33) or the 3-book test (n = 40) in Figure 2b ^2^. In Figure 2c the present results are compared to those obtained previously using the iPod stereo test in which the incremental depth judgement that was depended on its polarity (i.e., infront or behind) ^1^. The present results that do not depend on depth polarity are much more tightly distributed than the previous ones^1^ that did depend on depth polarity, indicating much less variability across the population when depth polarity is not involved.

**Figure 2.**
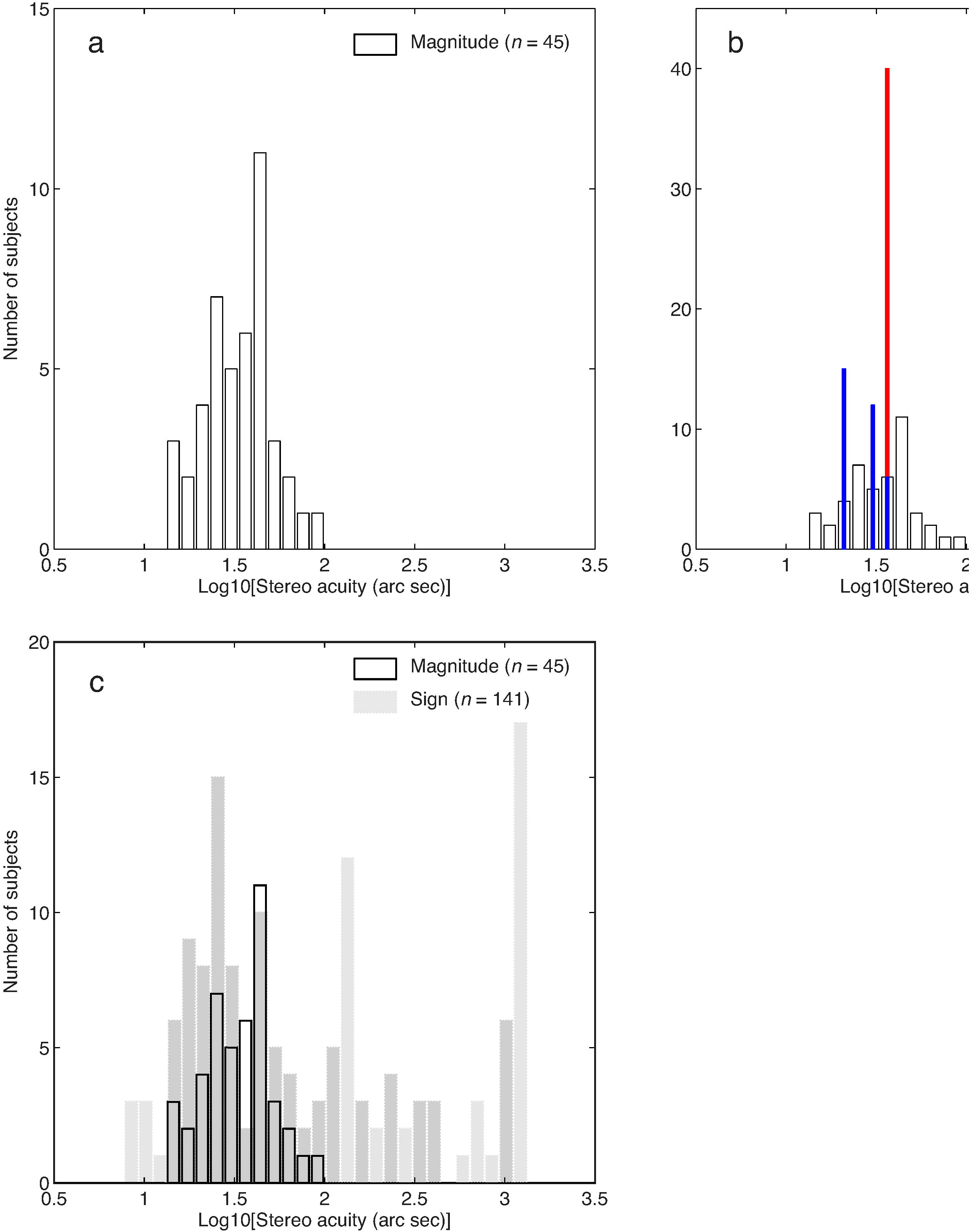
Histograms to show the stereo acuity results. (a) Stereo acuity results for 45 normal observers using a single polarity depth increment task (Pacman shape test); (b) The present results are compared with those of a previous study (shape defined by a single polarity depth increment) using the Randot preschool stereo test on individuals between the ages of 19-35 using the 4-book test (blue histograms) or the 3-book test (red histograms) ^2^; (c) the present results are compared to those obtained previously using the same approach but where the judgment involved stereo polarity (i.e., polarity-based increment task; grey histograms) ^1^.

## Discussion

Our results confirm that the detection of stereo-deficiencies within the normal population crucially depends on whether the measurement of stereopsis is dependent on the polarity of depth (i.e., a polarity-based increment) or not (a single polarity increment), confirming the earlier suggestions of several researchers ^1, 3–9^. As a consequence, it is a great pity that there is not currently a clinical test that can detect these deficiencies in the normal population and do it in a reliable way with an associated measure of variability. The current book tests (Randot and TNO but see^11^ for complete list) only measure depth detection for a fixed polarity (i.e., depth magnitude) and hence would miss the stereo-deficiencies reported here. The Frisby test (Burnell Co) does involve a depth polarity judgement but lacks an associated measure of variance, necessary for within-subject comparisons. It also has associated monocular cues^12^. The current iPod stereo test offers a number of important advantages over what is currently in use.

### Alternate explanations

*Could it be that the Randot test*^*2*^ *with its sharp edges can reveal finer stereo than stimuli with gaussian filtered edges*^*1*^ *and that polarity has nothing to do with the discrepancy?* No, other polarity-based tests with sharp edges^3^ highlight the same discrepancy. *Could there be an issue with the fusion of random dot stimuli that have gaussian blurred edges* ^*1*^? No, for the same reason outlined in the previous answer. Gaussian filtered edges are an effective method to avoid monocular cues that can be produced by sharply defined areas of depth within random dot arrays. *Could it be due to an increased positional uncertainty due to the Gaussian filtered edges leading to an uncertainty in the shape judgement?* No, Both the 2016 (Hess et al, 2016) and the present study used stimuli with gaussian filtered edges. The present study relied more on the form of the disparity, as the orientation in the pacman had to be detected. However, this study (depth magnitude measure), unlike the 2016 study (depth sign measure) did not show a wide variation in stereo acuity of normal observers. *Could it be that when stimuli of two depth polarities are displayed together, the visual system pools them and the problem highlighted is not one of polarity per se but anomalous depth pooling of simultaneous presented targets of opposite polarity?* ^9^, using figural instead of random dot stimuli, showed that adults (but not children) exhibit deficient depth discrimination on a supra-threshold task (for disparities above 288 seconds) only when opposite polarity stimuli are simultaneously presented and not when single polarity stimuli are presented. This suprathreshold measure is different from our threshold acuity measurements but the conclusions are broadly similar. They felt, for their suprathreshold measure, that anomalous disparity pooling might be responsible for two simultaneously presented depth surfaces. We don’t see this as such a good candidate explanation for our threshold measure because the stereo anomaly we report has been shown even when the two polarity surfaces are presented successively^3^. The role of the spatial layout of the stimuli (i.e., stimulus configuration, 2d grouping properties and overall scale of the stimulus array) or the exposure duration (down to 150ms) was shown to be unimportant in the Wilcox et al study.

### Proposed explanation

We have assumed that when a subject can detect a shape defined by a crossed disparity in the case of the Randot or other such clinical tests that they also have access to the fact that the surface appears in front of the page. This is an unwarranted assumption. Take the example of motion perception, a process often compared with stereopsis. The detection of a shape defined by a subset of dots moving in a particular direction does not necessarily require a knowledge of the direction of movement, it could be done simply by non-directional temporal information. A lesion in extra-striate cortex can abolish “motion perception” without affecting temporal change perception which can be accomplished in striate cortex^13, 14^. Our judgement of depth magnitude and its associated sign may be processed at different stages alone the cortical pathway and may be subject to different limitations. This is very much along the lines of a proposal made by Landers and Cormack^8^, who also reported a large percentage of stereo anomalous observers in the population for determining the sign but not the magnitude of the depth. They suggested that while there may be a continuous representation of disparity within the visual cortex, attentive readout of disparity may be a multicomponent serial process, having the following stages; starting from detection of disparite areas (local correlation), magnitude of disparity (present clinical stereo tests and the stimulus used in this present study), detection on the basis of disparity polarity ^1^) and finally discrimination of depth polarity. We propose that the percentage of normal subjects that exhibit anomalous stereopsis will simply depend on which of these stages is being targeted by the test; the higher the processing stage, the greater the percentage of stereo anomaly within the normal population.

### Clinical as well as real world relevance

Stereopsis is measured in the clinic for a number of reasons. First and foremost, as an indicator of the quality of binocular function. It is also used sometimes for occupational reasons as some careers require stereoscopic abilities, for example pilots, crane operators etc. Do the conclusions of this laboratory study invalidate the use of current stereo tests that are based solely on incremental depth judgements of a single polarity? The answer has to be yes. In normal binocular vision, it is implied that we can discriminate the “infront” as well as the “behind” depth of objects. An inability to do this constitutes an impairment. The majority of current stereo book tests only measure in front or crossed disparities and it is assumed that the same is true for behind or uncrossed disparities. That is simply not the case in up to 30% of the population. In terms of occupational needs, being able to discriminate the polarity of depth (infront vs behind) is obviously crucial. The next step in this research is to translate what has been shown in the laboratory using computer generated disparities to assess their real-world consequences.

